# Integrated Systems-Level Proteomics and Metabolomics Reveals the Diel Molecular Landscape of Diverse Kale Cultivars

**DOI:** 10.1101/2022.04.09.487754

**Authors:** Sabine Scandola, Devang Mehta, Brigo Castillo, Nicholas Boyce, R. Glen Uhrig

## Abstract

Kale are a group of diverse *Brassicaceae* species that are nutritious leafy greens consumed for their abundance of vitamins and micronutrients. Typified by their curly, serrated and/or wavy leaves, kale varieties have been primarily defined based on their leaf morphology and geographic origin, despite having complex genetic backgrounds. Kale is a very promising crop for vertical farming due to its high nutritional content; however, being a non-model organism, foundational, systems-level analyses of kale are lacking. Previous studies in kale have shown that time-of-day harvesting can affect its nutritional composition. Therefore, to gain a systems-level diel understanding of kale across its wide-ranging and diverse genetic landscape, we selected nine publicly available and commercially grown kale cultivars for growth under near-sunlight LED light conditions ideal for vertical farming. We then analyzed changes in morphology, growth and nutrition using a combination of phenomics, proteomics and metabolomics. As the diel molecular activities of plants drive their daily growth and development, ultimately determining their productivity as a crop, we harvested kale leaf tissue at both end-of-day (ED) and end-of-night (EN) time-points for all molecular analyses. Our results reveal that diel proteome and metabolome signatures divide the selected kale cultivars into two distinct groups, defined by their amino acid and sugar content, along with significant proteome differences involving carbon and nitrogen metabolism, mRNA splicing, protein translation and light harvesting. Together, our multi-cultivar, multi-omic analysis provides robust quantitative insights into the molecular underpinnings of the diel growth and development landscape of kale, significantly advancing our fundamental understanding of this nutritious leafy green super-food for next-generation horticulture / vertical farming applications.

## INTRODUCTION

*Brassica oleracea* and its diverse cultivar groups represent an important food crop for multiple populations across globe. These include seven major cultivar groups: cauliflower, collard greens, broccoli, kohlrabi, cabbage, brussels sprouts and kale. Together, these crops represented 70.1 million metric tonnes of production in 2019 (https://www.fao.org/faostat/en/#data/QCL). Kale, which encompass several leafy Brassicaceae species (*B. oleracea* and *B. napus*) (Reda et al. 2021), is often referred to as a ‘super-food’ (Samec et al. 2019) as it is rich in numerous antioxidants (carotenoids, flavonoids, glucosinolates) and essentials vitamins (A, K and C), minerals (calcium and iron), dietary fibers (Megias-Perez et al. 2020) and low molecular weight carbohydrates (Becerra-Moreno et al. 2014). Additionally, kale has notable cultivation advantages, including a wide-ranging temperature tolerance that guarantees year-round availability in most climates (Samec et al. 2019). Given these characteristics, kale represents a horticultural crop with the potential to be a source of essential nutrients for multiple global populations (Migliozzi et al. 2015).

To maximize growth and to execute developmental programs, plants require the precise timing of diel (daily) events. Correspondingly, diel events are coordinated by a combination of circadian and light responsive mechanisms, which play a major role in modulating the plant cell environment at all molecular levels (Mehta et al. 2021). For example, in the model plant and related Brassicaceae, *Arabidopsis thaliana* (*Arabidopsis*), it is estimated that the circadian clock controls the diel expression of 1/3 of all genes (Covington et al. 2008). Further, the importance of the circadian clock and diel biology in plants is emphasized by its central role in governing critical agronomic traits such as biomass, flowering time and disease resistance (Creux and Harmer 2019), suggesting diel biology should be a central facet of next-generation cropping systems (Hotta 2021; Steed et al. 2021). This is particularly key for kale, which has shown that time-of-day harvesting can affect its nutritional composition (Casajus et al. 2021; Francisco and Rodriguez 2021), indicating that time-of-day harvesting and postharvest storage is central to enhanced kale nutrition and shelf life when going to market (Reda et al. 2021). Currently our understanding of kale rests at the production-level, with the diel molecular mechanisms underpinning unique and beneficial morphological and nutritional differences between kale cultivars remaining largely unknown (Megias-Perez et al. 2020).

To date, systemic molecular analyses of kale cultivars *convar. acephala* (Jin et al. 2018; Liu et al. 2020; Liu et al. 2021) *var. sabellica* (Pongrac et al. 2019) and *B. napus var. pabularia* (Chiu et al. 2018) have been limited, with no studies examining multiple cultivars or quantifying diel molecular changes. In one transcriptomic study of *B. oleracea convar. acephala* cultivars from Asia, glucosinolate, carotenoid and phenylpropanoid biosynthetic pathways were highlighted as critical in defining the differences between green ‘manchoo collar’ and red ‘jeok seol’ cultivars (Jeon et al. 2018). To date however, no investigations of *B. oleracea var. palmifolia* have been performed, limiting our molecular knowledge of these widely produced and consumed kale cultivars. This set of molecular studies has however demonstrated that kale is a highly dynamic and diverse set of species that requires a systemic, multi-omics investigation of multiple kale cultivars in order to elucidate diel features of the molecular landscape that can be harnessed for increased production and nutrition.

Light spectra, intensity and photoperiod have each been shown to be important in kale cultivation as modulators of the kale metabolome (Carvalho et al. 2014). Light is also essential for plant growth and development as it is a primary entrainment mechanism of the circadian rhythm of plants (Xu et al. 2022). With the circadian clock and diel plant cell regulation governing numerous agronomic traits of interest, including: flowering time, growth and plant defense (Steed et al. 2021), elucidating the molecular landscape of kale across diverse genetics through a diel / circadian lens using quantitative, systems-level omics technologies, represents a critical endeavour for optimizing the growth, development and nutrient content of kale grown in controlled growth environments.

Therefore, using a combination of gas-chromatography mass spectrometry (GC-MS) and the latest data independent acquisition (DIA) quantitative proteomics workflow called BoxCarDIA (Mehta et al. 2022), we establish the diel metabolome and proteome landscapes of nine widely available, commercial produced kale cultivars grown in controlled growth environments under a natural photoperiod using near-sunlight conditions. These growth conditions, combined with multi-omics analyses obtained at end-of-day (ED) / zeitgeber 11 (ZT11), and end-of-night (EN) / zeitgeber 23 (ZT23) time-points, provides critical new insights into how diverse kale cultivars coordinate diel molecular events, while simultaneously generating a substantive proteome resource for further targeted experimentation. Through the development of this resource we lay a substantive foundation for uncovering the key proteins and cell processes underpinning critically important growth and development traits in kale, while also revealing the nutritional content and metabolic landscape of nine diverse kale cultivars.

## RESULTS

### Diversity in kale growth and morphology under horticultural LED light conditions

To compare the growth of nine publicly available and commercially grown kale cultivars (Table 1), we utilized spectral LED lighting conditions that can be implemented in LED driven horticultural growth systems. In addition, we further implemented twilight conditions at ED and EN to mimic a more natural growth environment (Figure 1A). We employed these parameters in order to comparatively evaluate the growth of all nine cultivars in a controlled growth environment setting that represents conditions comparable to those in future-forward vertical farming and horticultural facilities (Figure 1A). These conditions successfully grew all nine kale cultivars, with each cultivar presenting unique patterns of leaf size, shape and lobation consistent with previously observed morphologies. However interestingly, the leaf reddening typically observed in *B. napus var. pabularia* cultivars and the red *B. oleracea var. sabellica* ‘Scarlet’ (Figure 1B; (Carvalho and Folta 2014; Waterland et al. 2019) was not observed. The *B. oleracea var. sabellica* cultivars (Starbor (K8), Darkibor (K7), Winterbor (K2), Dwarf Curled Scotch (K3) and Scarlet (K9) presented an oval shaped, curly leaf phenotype, while *B. napus var. pabularia* cultivars (Red Russian (K5) and Red Ursa (K10) have a notably pronounced, serrated leaf morphology. Alternatively, the Italian cultivars *B. oleracea var. palmifolia* (Lacinato and Rainbow Lacinato) present a spear-like leaf phenotype (long and narrow), with a rough surface and darker green coloration relative to the other kale cultivars (Figure 1B). These leaf traits along with leaf size are correlated with each cultivar’s origins in Northern Europe, Russia and Italy, respectively (Table 1; Supplemental Figure 1). Lastly, we monitored leaf area over-time using a combination of time-course RGB imaging and PlantCV (https://plantcv.readthedocs.io/), which revealed kale cultivars to differ in their growth rates (Supplemental Figure 1). K10 and K2 varieties demonstrated the largest overall plant area, reaching an area of 13.4 to 12.6 cm^2^ at 24 days post-imbibition, while K8 and K5 exhibited the slowest growth rate reaching both a plant area of 8 cm^2^ at 24 days post-imbibition (Supplemental Figure 1). We also found that fresh weight (FW) is correlated with leaf area results, with K10 and K2 having the highest weight average of ∼ 4 g and K8 and K5 the lowest, with an average of 1 g and 0.8 g, respectively.

**Table 1:**
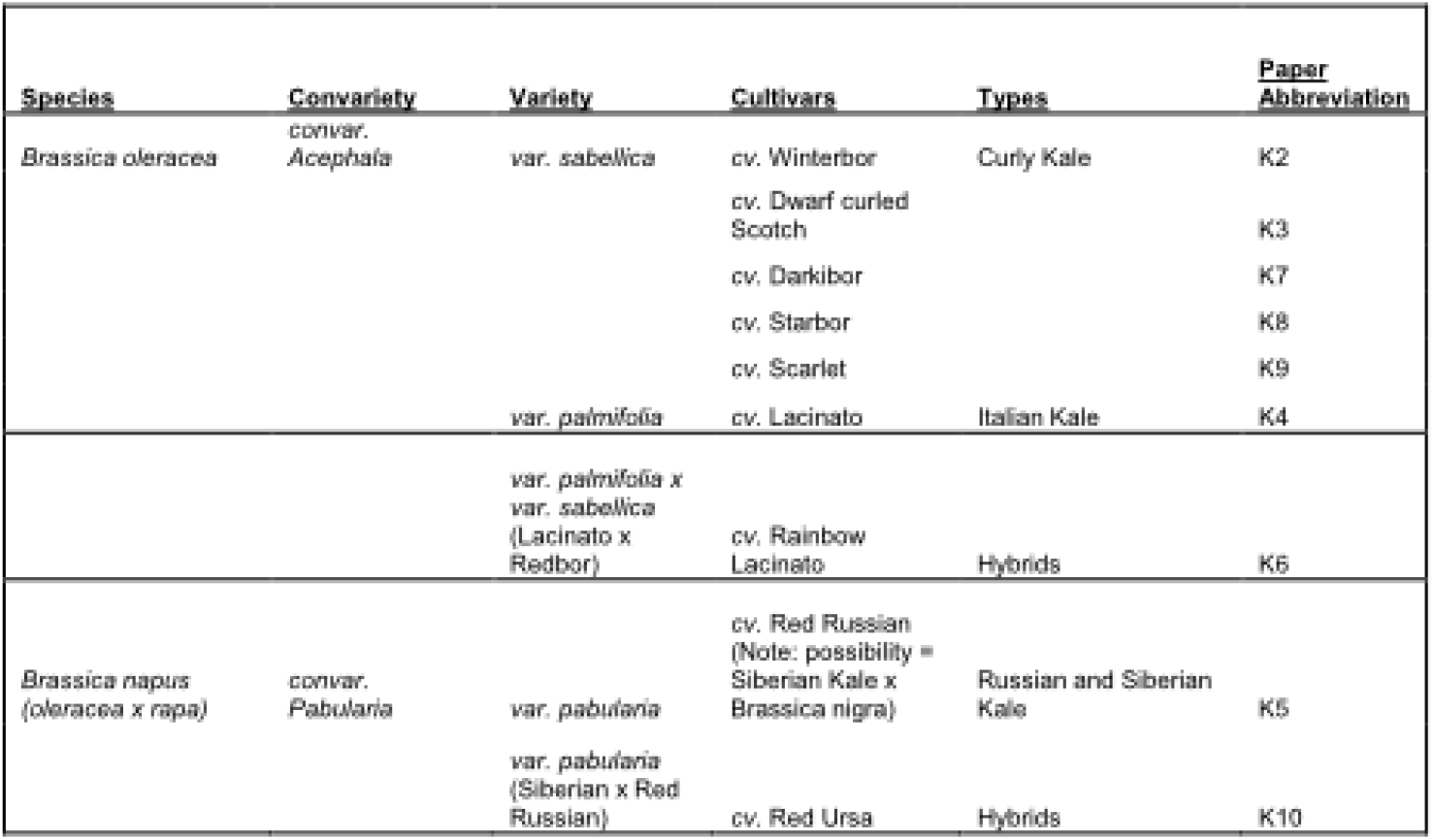
Name and classification of kale cultivars used in the study.

**Figure 1:**
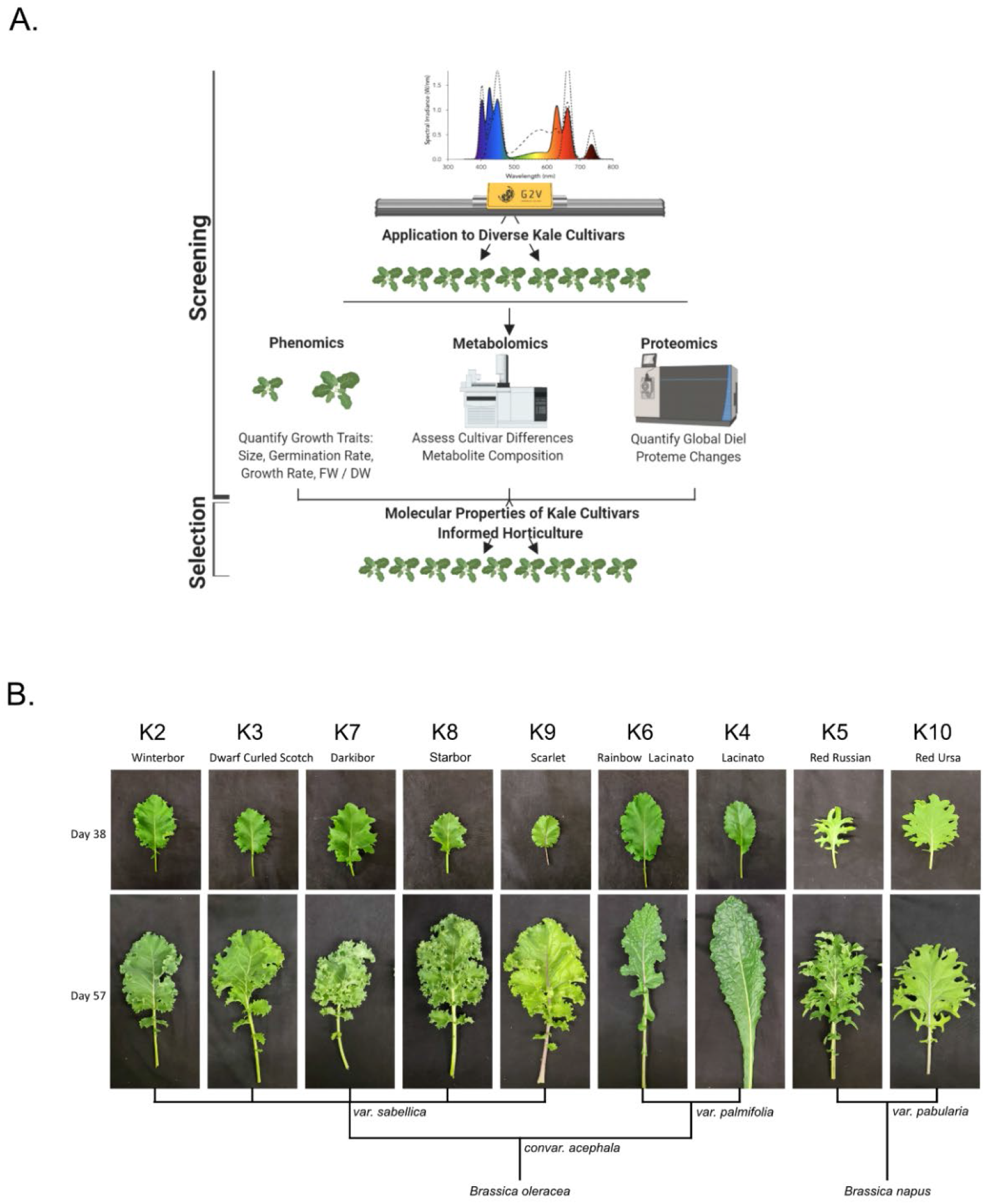
Experimental design and kale genetics phenotype. (A) Experimental workflow schematic. (B) Classification of kale cultivars based on leaf morphology and species.

To better characterize each cultivars physiological responses, we next measured relative chlorophyll content or Special Products Analysis (SPAD) and a variety of photosynthetic parameters (Phi2, PhiNO, PhiNPQ, LEF, ECSt, gH+ and vH+) using the PhotosynQ platform and the handheld MultispeQ device (Kuhlgert et al. 2016). Across the cultivars, relative chlorophyll amount was not significantly different except for the *var. palmifolia* cultivars K4 and K6, where SPAD was significantly higher (Supplemental Figure 1). This is consistent with the darker phenotype of the two *var. palmifolia* cultivars (Figure 1). Phi2, which measures photosystem II quantum yield, was only significantly higher in K5 compared to K10, while exhibiting no significant differences amongst other cultivars. Alternatively, PhiNO, which is a measurement of the electrons lost to non-regulated processes that can result in cellular damage, was significantly higher in K10, demonstrating that while K10 possesses one of the largest leaf areas, it is less effective at harvesting light energy. Interestingly, we find that linear electron flow (LEF), which estimates photosynthesis, exhibits a trend inversely proportional to FW, with large area cultivars possessing lower LEF and small cultivars possessing higher LEF. However, significant differences were only observed between cultivars K6 and K2.

Lastly, we estimated the energy generating capacity of each cultivar by measuring a series of parameters relating to ATP generation (Supplemental Figure 1). This included: ECSt, gH+ and vH+. ECSt, which describes the magnitude of the electrochromic shift, was higher in the K2 cultivar compared to K4, K8 and K9, suggesting better ATP production for increased growth outcomes while also aligning with their increased growth rate relative to other kale cultivars. Conversely, thylakoid proton conductivity (gH+), which describes steady state proton flux, was highest in the smallest cultivar K9, suggesting more efficient energy generation. Interestingly, despite differences in the ECSt and gH+, the initial rate of proton flux through ATP synthase (vH+) was constant across the cultivars. Taken together, these results indicate that photosynthetic parameters LEF and ECSt define important physiological differences in kale cultivars that likely contribute to observed differences in morphology and biomass.

### Diel metabolome analysis

As previous research in the related Brassicaceae *Arabidopsis* has demonstrated the importance of the ED and EN photoperiod transitions (Zeitgeber; ZT11-12 and ZT23-0; respectively) at the molecular-level (Krahmer et al. 2022; Uhrig et al. 2019), we next analyzed the metabolite content of each cultivar at ED (ZT11) and EN (ZT23). Here, we quantified diel changes in 40 metabolites across all nine cultivars, which can be grouped into 8 molecule classes: amino acids (12), organic acids (7), sugars (6), fatty acids (4), sterols (1), phenylpropanoid pathway metabolites (4) and vitamins (3) (Figure 2, Supplemental Figure 2; Supplemental Data 1). Euclidian distance hierarchal clustering further revealed that based on these 40 metabolites, the kale cultivars analyzed form 2 distinct groups based on their patterns of diel metabolite level fluctuations (Figure 2, Supplemental Data 1). This included a group consisting of Darkibor (K7), Starbor (K8), Red Russian (K5) and Red Ursa (10), which form Group I and Winterbor (K2), Dwarf Curled Scotch (K3), Scarlet (K9), Rainbow Lacinato (K6) and Lacinato (K4), which form Group II. This classification and grouping now allows us to establish more useful kale relationships based on metabolite landscapes (Supplemental Figure 2) rather than leaf morphology and geographic origins. Interestingly, Group I kale demonstrate more extensive leaf lobation and serration relative to their Group II counterparts, suggesting a correlation between leaf phenotype and diel metabolite changes. Further, hierarchal clustering analysis revealed two clusters of metabolites, whose relative change in diel abundance seem to be core to the differences between Group I and Group II (Figure 2). This includes: carbohydrates xylose, glucose and fructose; and amino acids aspartic acid, serine and iso-leucine, along with shikimate and α-Linolenic acid (Figure 2). Of the compounds that kale produces in larger quantities that are of direct nutritional importance, we detected vitamins niacin (vitamin B3), ascorbic acid (vitamin C) and phytol (vitamin E precursor) in addition to α-Linolenic acid (omega-3 fatty acid), with the majority of the kale cultivars possessing increased amounts of these vitamins at EN (Figure 2). Within Group II kale, we also see a sub-cluster consisting of K6 and K9 cultivars, which is largely defined by increased abundance of glyceric acid and glycine at ED.

**Figure 2:**
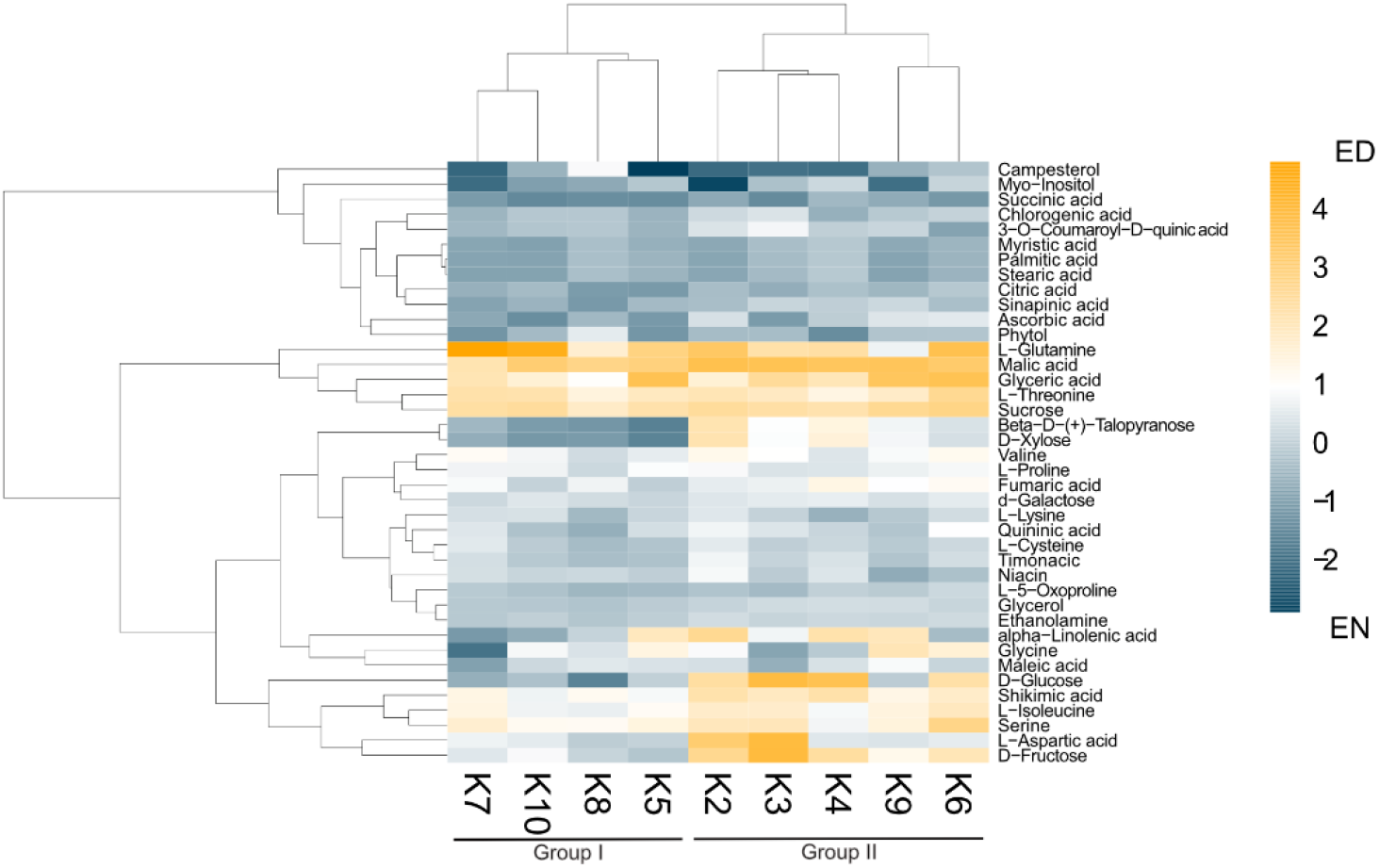
Diel changes in the kale metabolic landscape. Eucledian distance clustered heatmap of relative diel metabolite changes with each kale cultivar reveals two clusters of kale based on diel metabolic changes. Scale represents Log2 fold-change (FC); n=4.

### Diel Proteome Analysis

To further contextualize the metabolite defined Group I and Group II kale, we next performed quantitative proteomic analysis using an advanced data independent acquisition (DIA) workflow called BoxCarDIA which is aimed at better analyzing high complexity, high dynamic range plant samples (Mehta et al. 2022). With kale representing a diverse assemblage of plant species without specifically sequenced genomes, we performed our quantitative proteomic searches using the *B. oleracea var oleracea* proteome. Here, we were able to quantify a total of 2124 protein groups (Supplemental Data 2) across all nine cultivars. Of these, a total of 1734 protein groups exhibited a significant change in diel abundance (Bonferonni corrected *p-value* ≤ 0.05 and Log2FC ≥ 0.58) in at least one of the nine kale cultivars examined (Supplemental Data 2). Comparative quantification of the significantly changing proteins at ED and EN supported our metabolite-defined Group I and Group II clusters, with Group I kale possessing more proteins with a significant change in abundance at EN and Group II kale generally possessing more proteins changing at ED (Figure 3A).

**Figure 3:**
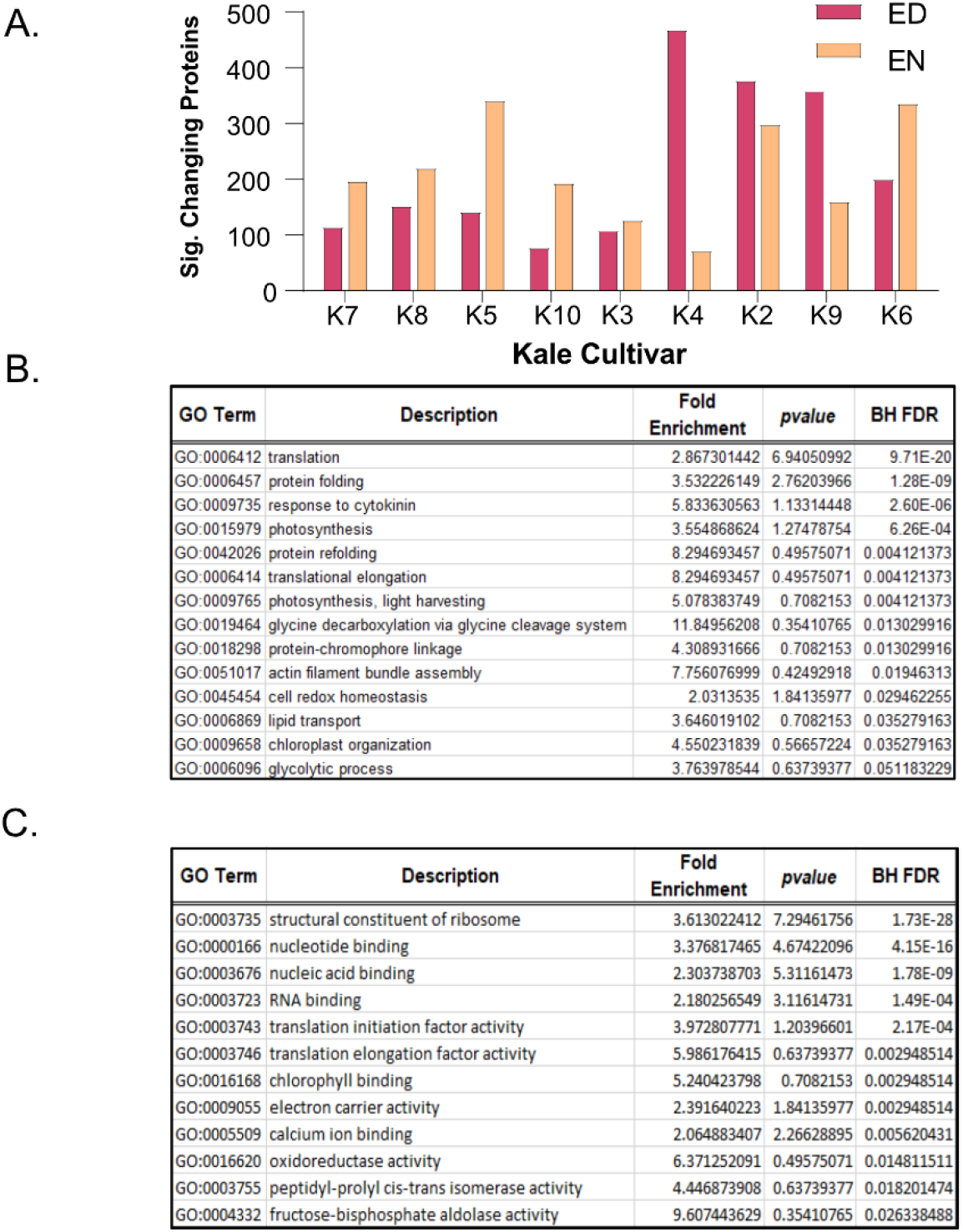
Diel changes in the kale proteome. Sampling diverse kale genetics provides a robust depiction of diel plant cell regulation in kale (n=4). (A) Number of proteins exhibiting diel changes in protein abundance within each kale cultivar examined at ED and EN. (B) Gene Ontology (GO) enrichment analysis of biological processes and (C) molecular function among all significantly changing proteins (Bonferonni corrected *q-value* ≤ 0.05; Log2FC ≥ 0.58 or ≤ -0.58).

Next, we analyzed all significantly changing proteins for enrichment of Gene Ontology (GO) terms relating to biological processes and molecular functions (BH corrected *p-value* ≤ 0.05). Here we found a significant enrichment of biological processes core to plant growth and development. These include significant enrichment of protein translation (GO0006412; GO:0006414), photosynthesis (GO:0009765; GO:0015979), cell redox homeostasis (GO:0045454), glycine metabolism (GO:0019464), lipid transport (GO:0006869) and primary metabolism (GO:0006096), amongst others (Figure 3B). Underpinning these biological processes was the significant enrichment of molecular functions related to translation initiation (GO:0003743) and elongation (GO:0003746), ribosome composition (GO:0003735), chlorophyll binding (GO:0016168) and oxidoreductase activity (GO:0016620), amongst other terms, which relate to protein translation, photosynthesis and cell redox homeostasis, respectively (Figure 3C).

To further elucidate when and where these protein-level changes differentially occur between Group I and II kale, and to increase our resolution of enriched biological processes, we performed an association network analysis using the knowledge database STRING-DB (https://string-db.org/). With *B. oleracea var oleracea* not possessing a STRING-DB dataset, we first identified orthologs from the related Brassicaceae *Arabidopsis* for all significantly changing proteins using UniProt (https://www.uniprot.org/). Correspondingly, we identified *Arabidopsis* gene identifiers for 80.4% (1395 / 1734) of the significantly changing proteins originally quantified (Supplemental Data 3). Using a highly stringent STRING-DB score of ≥ 0.9, we then mapped an association network for Group I and Group II kale. This revealed diel abundance changes in proteins related to RNA splicing, both cytosolic and plastidial translation, chlorophyll biosynthesis, chaperones, mitochondrial respiration and elements of carbon metabolism, with Group I exhibiting specific changes in the proteasome, protein secretion, fatty acid biosynthesis and methionine metabolism, while Group II maintained specific changes in the phagosome (Figure 4). STRING-DB analyses were further contextualized by subcellular localization data to elucidate where ED and EN changes manifest within the subcellular landscape (Figure 5). Group I predominantly exhibited changes at EN relating to proteins localized to the plastid, cytosol, mitochondria and extracellular compartments, while Group II exhibited most changes in similar compartments (plastid, cytosol and mitochondria), but at ED (Figure 5).

**Figure 4:**
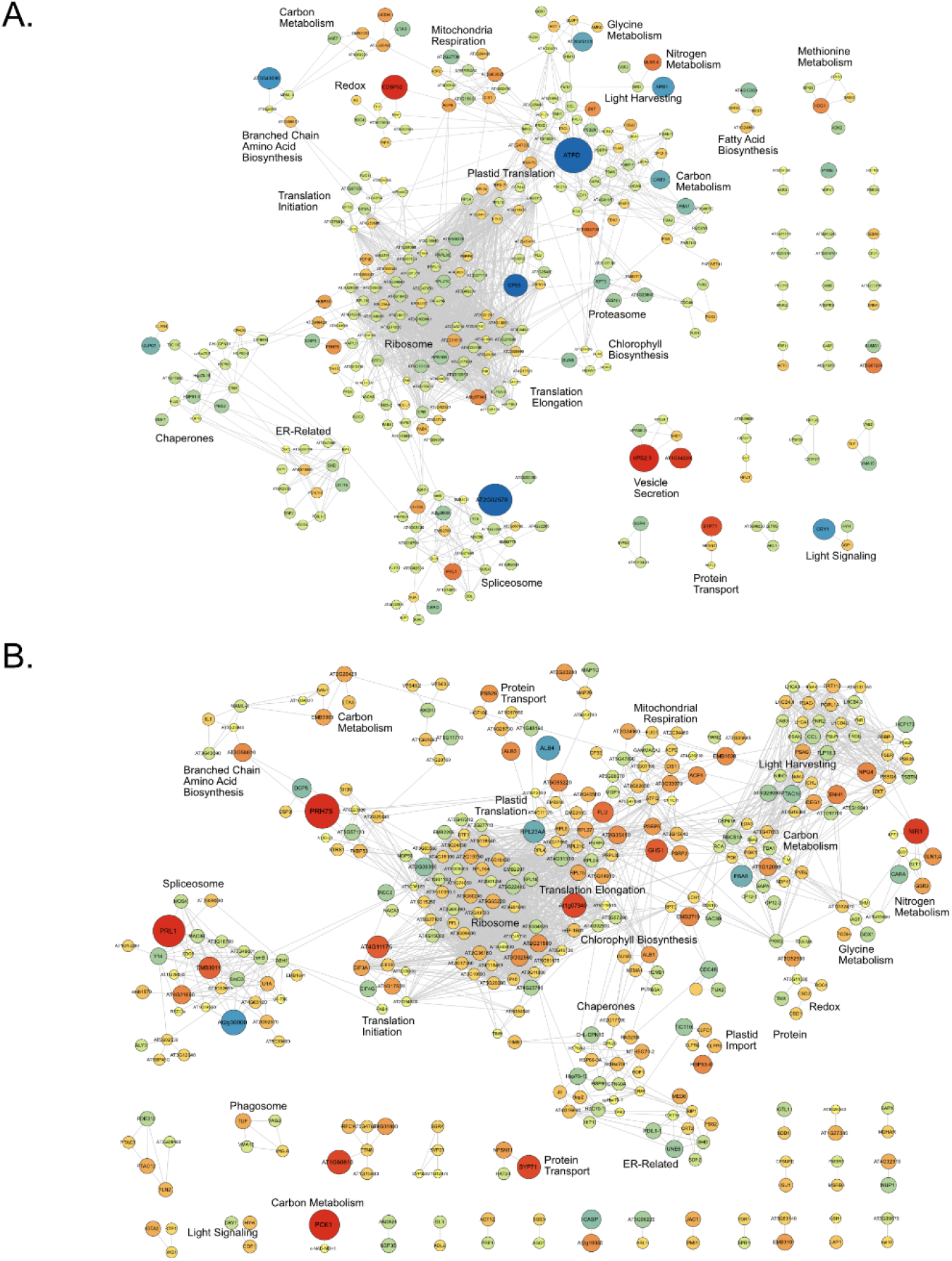
Association network analysis of the diel kale proteome. Association network analysis provides context for the networks of proteins exhibiting significant diel changes in protein abundance within Group I (A) and Group II (B) kale, respectively (https://string-db.org/). To maximize the analysis, *B. oleracea var. oleracea* gene identifiers were converted to *Arabidopsis* ortholog gene identifiers using UniProt (https://www.uniprot.org/). Proteins exhibiting significant changes in abundance are defined by a Bonferonni corrected *q-value* ≤ 0.05 and a Log2FC ≥ 0.58 or ≤ -0.58).

**Figure 5:**
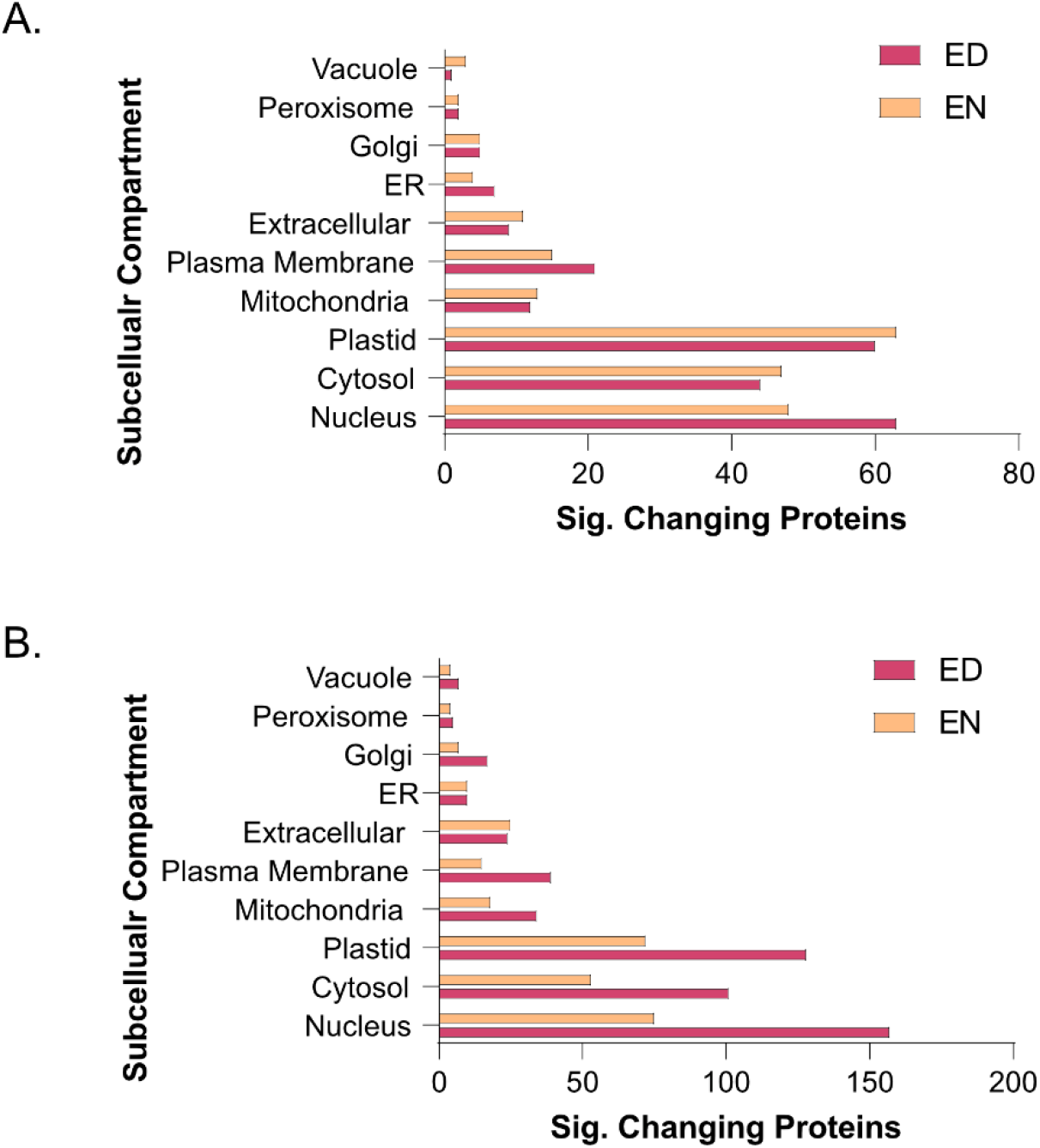
Subcellular localization analysis of the kale diel proteome. Subcellular localization analysis of Group I (A) and II (B) kale proteins exhibiting a significant change in diel abundance, respectively (https://suba.live/). To maximize this analysis, *B. oleracea var. oleracea* gene identifiers were converted to *Arabidopsis* ortholog gene identifiers using UniProt (https://www.uniprot.org/). Proteins exhibiting significant changes in abundance are defined by a Bonferonni corrected *q-value* ≤ 0.05 and a Log2FC ≥ 0.58 or ≤ -0.58).

## DISCUSSION

### Diverse kale cultivars form two distinct groups based on phenotypic and metabolic signatures

As the functional components of all biological systems and the defining elements of nutrition, the proteome and metabolome represent the most reliable means by which to elucidate differences between diverse, but related plant genetics. In the case of kale, it is the underlying molecular differences between kale cultivars that offer unique opportunities for targeted breeding and/or growth manipulation to enhance nutrition and/or biomass production. To date, differences between kale cultivars has been largely defined phenotypically through leaf morphologies such as coloration, size, shape, lobation and serration (Arias et al. 2021). A few studies have complemented this with transcriptomic analyses of individual cultivars (Arias et al. 2021; Chiu et al. 2018; Jeon et al. 2018; Jin et al. 2018), while others have undertaken metabolomics analyses (Chiu et al. 2018; Jeon et al. 2018). This has predominantly involved targeted metabolomics, examining single kale cultivars for changes in pigmentation (Redbor; *var. sabellica*; (Klopsch et al. 2019), flavonols (Winterbor; *var. sabellica*; (Neugart and Bumke-Vogt 2021; Neugart et al. 2014; Neugart et al. 2016) and fatty acids (Black Cabbage; *convar. acephala*; (Ayaz et al. 2006), with few studies having pursued global metabolite profiling of an individual kale cultivar (Nemzer et al. 2021). Correspondingly, our systems-level analysis of the leaf proteome and metabolome from nine kale cultivars in the same study using non-targeted GC-metabolomic and LC-proteomic mass spectrometry (MS), respectively, represents a substantial advancement in resolving the broader molecular landscape of kale, facilitating the creation of a critical resource for future targeted investigations.

Under our growth conditions no red/purple pigmentation in any kale cultivar was observed despite some of the cultivars examined (K9, K5 and K10) being known to have elevated anthocyanin production (Waterland et al. 2019). As a goal of our study was to define the diel molecular landscape of kale across diverse kale cultivars using near-sunlight conditions, our finding of no observable anthocyanin production was very informative. It suggests that LED light recipes deployed in controlled growth environments can be utilized to drive substantially different growth outcomes in the same kale variety. This aligns with previous studies, which revealed the application of UV light enhances the profile of flavonoids in kale (Neugart and Bumke-Vogt 2021; Neugart et al. 2014). Flavonoids are important molecule class in kale, as they represent the molecular precursors for the red/purple coloration some kale cultivars exhibit via anthocyanin (Liu et al. 2021a).

Unexpectedly, our analysis of the diel metabolome landscape across nine kale cultivars revealed two distinct kale groups based on their metabolite signatures that did not cluster along phylogenetic lines. Group I is comprised of both *B. oleracea* (K7 and K8) and *B. napus* (K5 and K10) cultivars, despite notable differences in ploidy between *B. oleracea* (diploid) and *B. napus* (polyploidy) (Gao et al. 2022). This suggests that higher level differences in genomic architecture do not seem to determine baseline growth traits in kale. Group I kale also maintained similarities in leaf lobation architecture, exhibiting a jagged and pronounced lobation morphology relative to the kale cultivars of Group II. Conversely, Group II kale consisted entirely of *B. oleracea* cultivars, which have substantially different and more variable lobation morphologies. The *var. palmifolia* cultivars K6 and K4, have a narrow leaf shape and almost no leaf lobation, while the *var. sabellica* cultivars K2, K3 and K9 have a round and wavy leaf shape with more leaf lobation. No specific photosynthesis or ATP production measurements were found to correlate with these metabolite-based groupings, likely due to the higher order nature of those processes relative to diel metabolite changes; however, Group II kale did possess more light harvesting / photosynthesis and mitochondrial respiration proteins exhibiting a diel change in their abundance relative to Group I. Using un-targeted metabolomics to define groups within a genus has also proven successful with other crops (*Cucumis melo*; (Moing et al. 2020). Here, a variable alignment between phylogeny and the metabolome was also found; paralleling our findings in kale. Taken together, our combined phenotypic and metabolomic definition of nine kale cultivars suggests that metabolomic fingerprinting provides a richer, more contextualized understanding of kale cultivars. Consequently, our findings provide a robust phenotypic and molecular resource for future fundamental research of this leafy-green super food, in addition to offering actionable information for vertical farming and horticultural kale producers.

### Underlying metabolic differences between kale cultivars offers opportunities for production systems

Upon comparing the diel metabolite profiles of our nine kale cultivars, we found a series of core compounds that are critical pre-requisites for the production of nutritionally valued specialized metabolites. Further, many of these metabolomic differences seemed to define Group I and II kale. These compounds include: carbohydrates (xylose, glucose and fructose), amino acids (serine, glycine, aspartic acid and iso-leucine) as well as shikimic acid and α-Linolenic acid. This analysis successfully defined the molecular potential of each kale cultivar, while also providing information for time-of-day kale harvesting in order to maximize its nutritional content. With the circadian clock and diel plant cell regulation highlighted as a critical consideration for next-generation agriculture (e.g. chronoculture (Steed et al. 2021), it is important that researchers and vertical farming / horticulture producers have reliable data resources generated using precision LED light systems.

#### Carbohydrates

Carbohydrates represent an important energy source for both plants and humans. Sugars, such as glucose and fructose, are a particularly important component of kale taste, which plays an important role in cultivar attractiveness, as kale is generally characterized as having a bitter and earthy flavor profile (Barker et al. 2022). Like other plants, kale possesses peak sucrose levels at ED, which in combination with transient leaf starch degradation, is vital for driving plant growth at night. At night, leaf starch is degraded to produce glucose, while sucrose is degraded to produce both glucose and fructose (Kim et al. 2017). Despite relatively consistent diel sucrose levels observed across both kale groups, we find a distinct diel pattern of glucose and fructose abundance between Group I and II kale. Group I kale possess more fructose at EN, suggesting they may have better cold-temperature tolerance as fructose enhances cold-induced oxidative stress adaptation by preserving homeostasis under low temperatures (Bogdanovic et al. 2008). This diel difference could also be explained by cultivars originating from northern latitudes (e.g. K5 and K10). Further, fructose may be involved in vitamin C biosynthesis through its conversion to fructose-6-phosphate, which also aids in cold tolerance (Akram et al. 2017), while also being an essential vitamin for human consumption. Diel changes in vitamin C leading to an accumulation at EN may offer to ease the transition to a high light environment come morning given its antioxidant properties (Paciolla et al. 2019). There is also a positive relationship between sucrose metabolism and anthocyanin production (Shi et al. 2014). This aligns with typically red cultivars K5 and K10, which both show enhanced diel accumulation of fructose and glucose at EN, while the typically green cultivars K4 and K6 of Group II kale possess maximal fructose levels at ED.

Additional carbohydrates xylose, myo-inositol and galactose were also present in different quantities across cultivars and throughout the day. Myo-inositol is involved in an array of biological processes such as being a precursor of inositol phosphates (Ips), hormones (auxin), translocation of mRNA into the cytosol, membrane biogenesis, light response germination and abiotic stress response (Munnik et al. 1998). Alternatively, xylose is a major component of the cell wall hemicellulose xylan, which provides plant cell resistance against enzymatic digestion and represents up to 35% of some wood compositions (Rennie and Scheller 2014). Xylose is derived from UDP-Glucose and is transported to Golgi where xylan is produced, which aligns with the large number of secretion related proteins (e.g. ER, Golgi and extracellular) we see significantly changing in both Group I and II kale. Xylose functions as a dietary fiber with prebiotic properties (Thavarajah et al. 2016). With Group I and II kale having contrasting diel xylose levels, the beneficial properties of xylose may offer value-added properties to a cultivar based on time-of-day harvesting.

#### Amino acids

Our analysis of the diel metabolome found that glycine, serine and aspartic acid have group specific differences in diel abundance. Both serine and glycine predominantly accumulated at EN and ED in Group I and II kale, respectively. Of these amino acids, serine in particular, functions as a key substrate for the biosynthesis of molecules critical for plant growth, including: amino acids glycine, methionine and cysteine (an essential amino acid for human nutrition), nitrogenous bases, proteins, phospholipids and sphingolipids (Stein and Granot 2019). Additionally, we find group specific differences in aspartic acid levels, which is also a precursor for several other amino acids, including: lysine, methionine, threonine and isoleucine (Han et al. 2021), with lysine representing an essential amino acid, whose content in plants is a key nutritional trait for crop improvement (Galili and Amir 2013). Amongst the kale cultivars examined however, we see lysine consistently produced at EN while we see iso-leucine demonstrating group specific diel abundance changes. Iso-leucine is one of three branched chain amino acids (BCAA; e.g. leucine, isoleucine and valine), but is the only BCAA not built from pyruvate (Joshi et al. 2010). It is also integral to plant defense as a conjugate of jasmonic acid (Armenta-Medina et al. 2021), in addition to being up-regulated in response to drought and cold (Joshi et al. 2010), suggesting that Group I and II kale may have notable differences on stress mitigation capacity.

#### Shikimic

Shikimic acid is a precursor of the essential aromatic amino acids tyrosine, phenylalanine and tryptophan and therefore is an important precursor for the phenylpropanoid biosynthetic pathway, which is responsible for the production of a large number of nutritionally valued specialized metabolites (Dong and Lin 2021). Production of shikimic acid can consume upwards of 30% of the fixed carbon, feeding the production of vitamins K1, B3 (folate), E (tocopherols), in addition to flavonoids, anthocyanins and lignin (Tohge et al. 2013). The shikimic acid pathway is also involved in the color patterning seen in ornamental kale through modulation of anthocyanin content, which is a key trait in kale as a source of antioxidants (Liu et al. 2021).

#### α-Linolenic acid

Kale is known to be rich in α-Linolenic acid, however, the differences in abundance between cultivars and the diel production landscape of α-Linolenic acid have not previously been assessed. α-Linolenic acid is a polyunsaturated Omega 3 fatty acid, which is an essential component of a healthy diet (Nemzer et al. 2021), as omega 3 fatty acids are known to decrease the risk of heart disease and lower the blood pressure (Shahidi and Ambigaipalan 2018). Importantly, α-Linolenic acid also forms the basis of cell membrane components as a precursor of phosphoglycerolipids, cutin and waxes (He and Ding 2020). With α-Linolenic acid contributing broadly to many of the important nutritional properties of kale and our results demonstrating it maintains diel changes in abundance, along with differences in abundance between cultivars, α-Linolenic acid offers an array of opportunities for further development of cultivar specific light recipes to maximize its production.

### Differences in the diel proteomes of Group I and II kale indicate they are defined by core elements of plant cell regulation

Elucidating the specific molecular components that underpin the observed diel metabolomic changes and growth characteristics of Group I and II kale revealed a number of plant cell processes that are diel regulated at the protein-level. In particular, we find diel proteome changes related to metabolism, RNA processing, protein translation and light harvesting. From a metabolic perspective, we find diel changes in: carbon, nitrogen, glycine, methionine, BCAA and fatty acid metabolism, while for RNA processing and protein translation, we find: mRNA splicing, cytosolic and plastidial protein translation along with a number of chaperones to exhibit significant diel changes. We also observed differences involving numerous light harvesting and signaling proteins, along with differences in chlorophyll biosynthetic enzymes, which may be directly related to Group specific productivity differences. At the highest level, our quantitative proteomic analysis resolved Group I kale cultivars to possess significant changes in their diel proteome at EN, while Group II kale exhibit more significant proteome-level changes ED. We also find that many significantly changing metabolic proteins are directly connected to observed diel metabolome changes, reinforcing our multi-omics approach. Further, significant changes in numerous other, non-metabolic proteins central to proper plant cell regulation demonstrate how quantitative proteome analysis allows us to map how the diel molecular landscape of kale changes in order to identify how kale breeders and producers can better realize each cultivars genetic potential.

#### Metabolism

Significant changes in the diel proteome related to plant metabolism were wide-ranging, aligning with the metabolite-determined kale groupings. Enzymes involved in carbon and nitrogen metabolism exhibiting a significant diel change in abundance demonstrated coherent time-of-day changes between both kale groups, consistent with their central roles in plant growth and development. Conversely, amino acid metabolic enzymes aligned with the metabolite-defined kale groups, specifically BCAA-related enzymes and glycine metabolic enzymes, likely relating to the specialized metabolites produced by kale that are derived from these amino acids.

From a carbon metabolism perspective, diel changes centered around two protein clusters comprised of mitochondrial enzymes isocitrate dehydrogenase (cICDH), ATP citrate lyase and components of the pyruvate dehydrogenase complex, along with key primary metabolic enzymes fructose-bisphosphate aldolase (FBA), glyceraldehyde 3-phosphate (GAPA) and phosphoglycerate kinase (PGK). With acetyl-CoA representing a critical metabolite involved in multiple biosynthetic pathways, including fatty acid biosynthesis (Xing and Poirier 2012), it is perhaps not surprising that cICDH and ATP citrate lyase enzymes exhibit consistent diel abundance changes in both Group I and II kale. Similarly, FBA, GAPA and PGK enzymes consistently exhibited significant diel abundance changes across both Group I and II, offering a consistent benchmark for our systems-level proteome analysis. Intriguingly, different isoforms of PGK and FBA were found to possess significant changes in diel abundance between Group I and II kale. In the related Brassicaceae *Arabidopsis*, the three encoded FBAs were found to have essential roles in plant metabolism, but with tissue specific expression patterns (Carrera et al. 2021). Here however, it seems that Group I and II kale each utilize a different subset of FBA isozymes, as we analyzed the same leaf tissues across cultivars. Unlike *Arabidopsis*, the polyploid *B. napus* has been shown to possess twenty-two FBAs, which possess diverse developmental expression patterns across multiple cellular compartments (Zhao et al. 2019). Further, it was also interesting that despite fructose and glucose representing two of the group defining metabolites in our study, no specific protein groups related to the generation of fructose and glucose such as starch degradation enzymes, sucrose synthase or invertase, were found to be significantly changing in our proteomics data. This perhaps indicates that changes in these enzymes, such as their activity are driven by other regulatory mechanisms such as reversible protein phosphorylation, rather than changes in abundance (Hardin et al. 2004).

We also quantified changes in nitrogen assimilating enzymes in both Group I and II kale. Group I kale cultivars possess substantially larger diel changes in nitrite reductase (NiR) levels at ED relative to Group II, while glutamine synthetase isoform 1,4 (GLN1,4), which converts glutamate to glutamine as part of nitrogen assimilation for transport from roots to shoots, exhibited a consistent and significant change in abundance at ED in both Group I and II kale. This aligns with our metabolite data, which finds glutamine levels to peak in nearly all nine kale cultivars at ED. In the model Brassicaceae *Arabidopsis*, it is well known that nitrogen metabolism is a diel regulated process (Flis et al. 2019), with peak transcript and protein abundance of nitrate reductase enzymes occurring early in the day, however here, it seems that the precise time-of-day coordination of these events differs between kale cultivars.

Connecting to primary metabolism, our proteomics data also resolved extensive proteome changes involving various facets of amino acid biosynthesis. Unlike carbon and nitrogen metabolism, amino acid metabolism closely parallels our metabolite data that defined Group I and II kale. In particular, we find significant differences in the diel abundance of enzymes related to glycine and BCAA (e.g. iso-leucine) metabolism, with glycine being directly connected to serine production, which we also observe having diel differences between Group I and II kale (Schulze et al. 2016). We also find time-of-day differences in iso-leucine production, which is produced down-stream of aspartate and possesses group-specific time-of-day production differences. Interestingly, despite aspartate fueling lysine and methionine production, we only find Group I kale possessing a significant change in methionine biosynthetic enzymes. These aligned diel differences in both the proteome and metabolome relating to amino acids makes a clear case for genetics-based time-of-day harvesting of kale for maximal nutritional content.

#### mRNA Processing & Protein Translation

RNA splicing and protein translation represent two cellular processes carried out by multi-subunit protein complexes; the spliceosome and the ribosome, respectively. However, much remains to be resolved as to how each of these complexes are regulated in plants at the protein level. Previous diel proteome analyses have found both the spliceosome and ribosome to be dynamically regulated at the protein-level by changes in both abundance and phosphorylation (Uhrig et al. 2021). However, resolution of species-specific differences in diel abundance has not previously been resolved. Here we see Group I and II kale defined by smaller abundance changes in mRNA processing and protein translation machinery at EN in Group I kale, coupled with larger abundance changes at ED in Group II kale. Correspondingly, a large abundance of chaperone proteins changing in a similar pattern at the same time-points is observed, likely aiding in effective protein production. Surprisingly, we also see a large abundance of plastidial translational machinery in both Group I and II kale, which parallel the diel changes in abundance found in their cytosolic counterparts, suggesting concerted coordination of global protein production in each kale group. Currently our understanding of diel changes in mRNA splicing and protein translation have largely been defined by transcriptomic sequencing technologies. In particular, use of polysome loading or RiboSeq as a proxy for protein translation, which in *Arabidopsis* is suggested to negatively correlate with biomass (Ishihara et al. 2017). Similarly, RNAseq profiling of *Arabidopsis* over a 24 h photoperiod have found diel changes in mRNA spliceforms (Romanowski et al. 2020), however, in both areas of research, direct diel assessment of spliceosome or ribosome complex composition and regulation at the protein-level has remained undefined. In light of the work performed in the related Brassicaceae *Arabidopsis*, our findings here suggest that there are systemic differences between the kale groups in the timing of growth as it relates to protein translation that could be utilized to enhance the productivity kale through the precise adjustment of growth conditions.

#### Light Harvesting and Signaling

In both Group I and II kale we see extensive diel changes in the light harvesting and photosynthetic machinery, with no specific ED or EN changes in either group. The largest changes observed involved the chloroplast ATP synthase delta subunit (ATPD) at EN in Group I kale along with enzymes ENHANCER OF SOS3-1 (ENH1) and NONPHOTOCHEMICAL QUENCHING 4 (NPQ4) at ED in Group II kale. Chloroplast ATP synthase is a critical driver of ATP production in plants in the light, with *Arabidopsis* plants lacking ATPD presenting a lethal phenotype due to the destabilization of the ATP synthase complex (Maiwald et al. 2003). Alternatively, ENH1 is required to main redox balance (Zhu et al. 2007) and NPQ1 is involved in non-photochemical quenching in the presence of excess light energy, which led to increased growth outcomes in tobacco when present in higher abundance (Kromdijk et al. 2016). Interestingly, in both Group I and II kale we also observe a small, but important network of proteins comprised of CRYPTOCHROME 1 (CRY1), ELONGATED HYPOCOTYL 5 HOMOLOG (HYH) and CONSTITUTIVE PHOTOMORPHOGENIC 1 (COP1), with CRY1 more abundant at EN in Group I kale. In response to blue light, CRY1 inhibits the degradation of the HY5 transcription factor by COP1 (Wang et al. 2018). Although there is a more limited understanding of HYH, HY5, an HYH ortholog, is a regulator of light-mediated transcription in plants, controlling a wide range of plant cell processes related to growth and development that are of importance to kale production (Xiao et al. 2021). CRY1 is also connected to the circadian clock through detection of blue light fluence (Somers et al. 1998; Sanchez et al. 2020), indicating potential higher-order regulation of timed-metabolism in Group I versus Group II kale. This has particularly intriguing chronoculture implications for Group I kale production in controlled growth environment settings given its larger diel abundance changes.

## SUMMARY

Our integrated, systems-level analysis of nine diverse, commercially produced and readily consumed kale cultivars has substantially advanced our understanding of kale from the phenotypic-level to the underlying molecular-level. In doing this, our dataset reveals new information about the molecular landscapes of these kale cultivars when grown under standardized controlled growth environment conditions, providing new opportunities for vertical farming and/or horticultural growth of kale. Our systems-level analysis has defined diel differences in the molecular landscapes underpinning these diverse kale genetics, elucidating information for time-of-day harvesting considerations to ensure maximal nutritional content. Variations in the diel molecular landscapes of different cultivars or plant accessions have been previously observed in *Arabidopsis* (Rees et al. 2021), tomato (Muller et al. 2016) and soybean (Greenham et al. 2017), with explanations for these differences being related to environmental stimuli and/or geography (seasonal impacts and latitude) (Rees et al. 2021; Ruts et al. 2012; Steppe et al. 2015). It is also interesting, that many of those differences have been linked to diel plant biology. Mechanistically, we observe differences in proteins linked to light perception and the circadian clock (e.g. CRY1), while simultaneously finding group-specific differences in multiple metabolites connected to the circadian clock (Cervela-Cardona et al. 2021; Haydon et al. 2013; Scandola et al. 2022). Overall, our endeavor to define the underlying molecular landscape of diverse, commercially grown kale cultivars has opened up extraordinary opportunities for horticultural production activities moving forward. Further, our results suggest that combined use of phenomics, proteomics and metabolomics represents a robust paradigm for characterizing non-model horticultural crops of diverse genetic backgrounds.

## MATERIALS AND METHODS

### Growth conditions

Nine commercially grown cultivars of kale were purchased from OCS Seeds (Table 1; https://www.oscseeds.com/) and West Coast Seeds (https://www.westcoastseeds.com/) and grown for the study. These included: *B. oleracea var. sabellica* cultivars Winterbor (K2), Dwarf Curled Scotch (K3), Darkibor (K7), Starbor (K8), Scarlet (K9), *B. oleracea var. palmifolia* cultivars Lacinato (K4), Rainbow Lacinato (K6) and *B. napus var. pabularia* cultivars Red Russian (K5) and Red Ursa (K10). Seeds were sterilized in 70% ethanol for 2 min followed by a 70% (v/v) bleach (Chlorox 7.5%) treatment for 7 min and 3 washes with distilled water. The seeds were then grown on ½ MS media containing 1% (w/v) sucrose and 7 g/L of agar at pH 5.8. The seeds were cold treated 3 days at 4°C in the dark and exposed to light for a week before being transferred to soil (Sun Gro®, Sunshine Mix® #1). At 29 days post-sterilization, entire plants were collected for GC-MS and LC-MS analysis. Growth chambers were equipped with a programmable Perihelion LED fixture (G2V Optics Inc; https://g2voptics.com/) and lined with Reflectix ® to ensure a good light distribution. Kales were grown under a 12h light and 12h dark regimen and a temperature of 21°C during the day and 19°C at night.

### Metabolite Extraction and GC-MS Analysis

#### Metabolite Extraction and Data Acquisition

Metabolite extraction and preparation were performed with modifications as previously described (Liu et al. 2016). Tissue was harvested and directly flash frozen in liquid nitrogen (n = 4; each biological replicate consists of a pool of 3 plants). Sample of 100 mg (+/-1 mg) of pulverized tissue were prepared and homogenized in 700 µl of iced-cold methanol (80% v/v). In each sample, 25 µl of ribitol at 0.4 mg.ml^-1^ in water were added as internal standard. Samples were incubated 2 h at 4 °C with shaking and then 15 min at 70°C at 850 rpm in a Thermomixer. Tubes were centrifuged 30 min at 12000 rpm and the supernatants were transferred in new tubes. Polar and non-polar phases were separated by the addition of 700 µl of water and 350 µl of chloroform, then vortexed thoroughly and centrifuged for 15 min at 5000 rpm. The upper methanol/ water phase (150 µl) was transferred to a new tube and dry in a vacuum centrifuge at RT. Samples were derivatized with 100 μl of methoxamine hydrochloride-HCl (20 mg.ml^-1^ in pyridine) for 90 min at 30°C at 850 rpm in thermomixer and followed by incubation with 100 µL of N,O-bis(trimethylsilyl)trifluoroacetamide (BSTFA) at 80°C during 30 min with shaking at 850 rpm in thermomixer. Finally, samples were injected in split less mode and analyzed using a 7890A gas chromatograph coupled to a 5975C quadrupole mass detector (Agilent Technologies, Palo Alto, CA, USA). In the same manner, 1 µl of retention time standard mixture Supelco C7–C40 saturated alkanes (1,000 µg.ml^-1^ of each component in hexane) diluted 100 fold (10 µg.ml^-1^ final concentration) was injected and analyzed. Alkanes were dissolved in pyridine at 0.22 mg.ml^-1^ final concentration. Chromatic separation was done with a DB-5MS capillary column (30 m × 0.25 mm × 0.25 µm; Agilent J&W Scientific, Folsom, CA, USA). Inlet temperature was set at 280°C. Initial GC Oven temperature was set to 80°C and held for 2 min after injection then GC oven temperature was raised to 300°C at 7°C min^-1^, and finally held at 300°C for 10 min. Injection and ion source temperatures were adjusted to 300°C and 200°C, respectively with a solvent delay of 5 min. The carrier gas (Helium) flow rate was set to 1 ml.min^−1^. The detector was operated in EI mode at 70 eV and in full scan mode (m/z 33–600).

#### Metabolites Data Analysis

Compounds were identified by mass spectral and retention time index matching to the mass spectra of the National Institute of Standards and Technology library (NIST20, https://www.nist.gov/) and the Golm Metabolome Database (GMD, http://gmd.mpimp-golm.mpg.de/). Metabolite quantification was performed using MassHunter Software from Agilent. Peaks were deconvoluted and integrated and were normalized to the internal standard ribitol and by the sample weight.

### Protein extraction and nanoflow LC-MS analysis

#### Protein Extraction and Data Acquisition

Kale leaf tissue was quick frozen and ground to a fine powder under liquid N_2_ using a mortar and pestle and aliquoted into 400 mg fractions (n = 4; each biological replicate consists of a pool of 3 plants). Samples were then extracted at a 1:2 (w/v) ratio with a solution of 50 mM HEPES-KOH pH 8.0, 50 mM NaCl, and 4% (w/v) SDS. This included vortexing, followed by incubation at 95°C in an Eppendorf microtube table-top shaking incubator shaking at 1100 RPM for 15 mins. This was then followed by an additional 15 mins of shaking at room temperature. All samples were clarified at 20,000 x g for 5 min at room temperature, with the supernatant retained in fresh Eppendorf microtubes. Sample protein concentrations were measured by bicinchoninic acid (BCA) assay (23225; ThermoScientific), followed by reduction with 10 mM dithiothreitol (DTT) at 95°C for 5 mins. Samples were then cooled and alkylated with 30 mM iodoacetamide (IA) for 30 min in the dark without shaking at room temperature. Subsequently, 10 mM DTT was added to each sample, followed by a quick vortex, and incubation for 10 min at room temperature without shaking. Total proteome peptide pools were generated by sample digestion overnight with 1:100 sequencing grade trypsin (V5113; Promega). Generated peptide pools were quantified by Nanodrop, followed by acidification with formic acid to a final concentration of 5% (v/v) and then dried by vacuum centrifugation. Peptides were then desalted using ZipTip C18 pipette tips (ZTC18S960; Millipore) as previously described ^7^, dried and dissolved in 3.0% ACN/0.1% FA prior to MS analysis. Digested samples were then analysed using a Fusion Lumos Tribrid Orbitrap mass spectrometer (Thermo Scientific) in a data independent acquisition (DIA) mode using the BoxCarDIA method as previously descried ^31^. Dissolved peptides (1 µg) were injected using an Easy-nLC 1200 system (LC140; ThermoScientific) and separated on a 50 cm Easy-Spray PepMap C18 Column (ES903; ThermoScientific). Liquid chromatography and BoxCar DIA acquisition was performed as previously described without deviation (Mehta et al. 2022).

#### Proteomic Data Analysis

All acquired BoxCar DIA data was analyzed in a library-free DIA approach using Spectronaut v14 (Biognosys AG) using default settings. Data were searched using the *B. oleracea* var *oleracea* proteome (Uniprot: https://www.uniprot.org/ containing 58,545 proteins). Default search parameters for proteome quantification were used, with specific search parameters including: a protein, peptide and PSM FDR of 1%, trypsin digestion with 1 missed cleavage, fixed modification including carbamidomethylation of cysteine residues and variable modifications including methionine oxidation. Data was Log2 transformed and globally normalized by median subtraction with significantly changing differentially abundant proteins determined and corrected for multiple comparisons (Bonferroni-corrected *p-value* ≤ 0.05; q-value).

### Bioinformatics

Gene Ontology enrichment analyses were performed using the Database for Annotation, Visualization and Integrated Discovery (DAVID; v 6.8; https://david.ncifcrf.gov/home.jsp). Significance was determined using Benjamini-Hochberg (BH) corrected *p-value* ≤ 0.05. Conversion of *B. oleracea var oleracea* gene identifiers to *Arabidopsis* gene identifiers for STRING association network analysis and SUBA4 subcellular localization information retrieval was performed using UniProt (https://www.uniprot.org/). STRING association network analyses were performed in Cytoscape v3.9.0 (https://cytoscape.org/) using the String DB plugin stringApp, all datatypes and a minimum correlation coefficient setting of 0.9. Predicted subcellular localization information was obtained using SUBA4 and the consensus subcellular localization predictor SUBAcon (https://suba.live/). The Eucledian distance *pheatmap* clustered heatmap analysis of diel metabolites was performed in R (3.6.1 R Core Team, 2019). Additionally figures were assembled using Affinity Designer software (v1.9.1.179; https://affinity.serif.com/en-us/designer/)

## DATA AVAILABILITY

All raw data files, mass spectrometry parameters, and Spectronaut search settings have been uploaded to ProteomeXchange (http://www.proteomexchange.org/) via the PRoteomics IDEntification Database (PRIDE; https://www.ebi.ac.uk/pride/). Project Accession: PXD031780.

## AUTHOR CONTRIBUTIONS

Plant growth, phenotyping and harvesting was performed by SS, NB and BC. Metabolite data acquisition and analyses were performed by SS. Proteomics data acquisition and analysis was performed by DM, SS and RGU. The manuscript was written by SS and RGU, with editorial input from DM.

## ACKNOWLEDMENTS

We would like to thank G2V Optics Inc. for providing their programmable LED lighting system. Thanks to Jack Moore and The Alberta Proteomics and Mass Spectrometry Facility (APM) for providing proteomic support for this work. Funding for this work was generously provided by the Natural Sciences and Engineering Research Council of Canada (NSERC) and Alberta Innovates through their Alliance and Campus Alberta Small Business Engagement (CASBE) funding schemes, respectively. This work was also supported by the Canada Foundation for Innovation (CFI).

**SUPPLEMENTAL FIGURES**

**Supplemental Figure 1: Kale morphology, growth and photosynthetic parameters comparison**. (A) Side and top pictures of kale cultivars at 35 days post imbibition (DPI). (B and C) Fresh weight at 35 DPI and means of kale area measured with PlantCV over the course of a day at 24 DPI (n≥4). (D to K) Relative chlorophyll content (SPAD) and photosynthetic parameters measured with MultispeQ at 38 DPI (n=10). Means with no letter in common are significantly different (Tukey’s multiple comparison test; p<0.05). Errors bars indicate ± SEM.

**Supplemental Figure 2: Diel changes in the kale metabolic landscape**. Cartoon schematic of mapped diel metabolic changes observed in kale cultivars by GC-MS.

**SUPPLEMENTAL DATA**

**Supplemental Data 1: Gas chromatography metabolite data Supplemental Data 2: Liquid chromatography proteomic data**

**Supplemental Data 3: Uniprot gene identifier conversion from *Brassica oleracea* to *Arabidopsis***

**Supplemental Data 4: PlantCV data and data processing Supplemental Data 5: MultisynQ phenotype data**

