## Supplemental Figure 1 for "Integrated Systems-Level Proteomics and Metabolomics Reveals the Diel Molecular Landscape of Diverse Kale Cultivars"

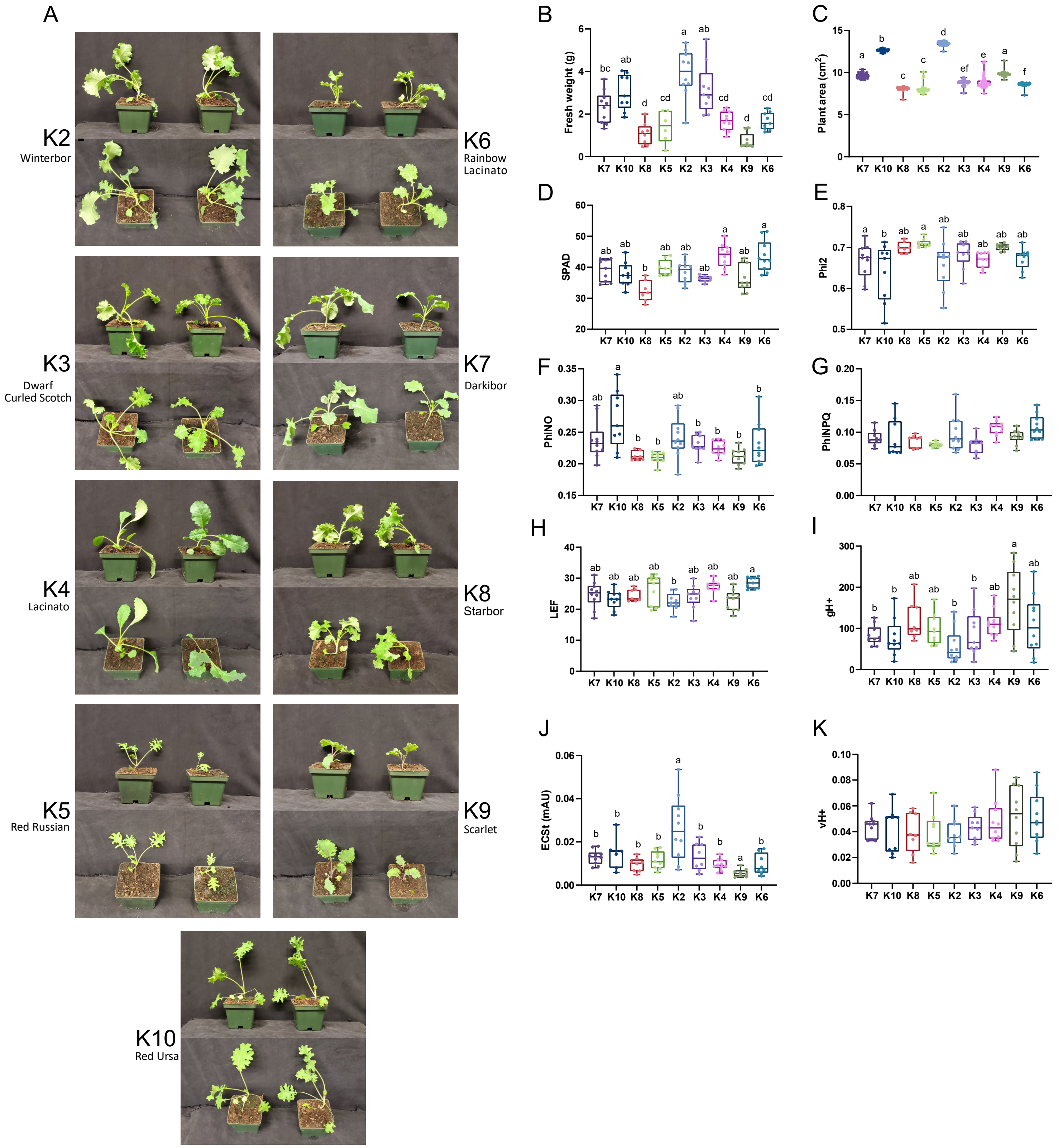


**Supplemental Figure 1: Kale morphology, growth and photosynthetic parameters comparison.** (A) Side and top pictures of kale cultivars at 35 days post imbibition (DPI). (B and C) Fresh weight at 35 DPI and means of kale area measured with PlantCV over the course of a day at 24 DPI ($n \geq 4$). (D to K) Relative chlorophyll content (SPAD) and photosynthetic parameters measured with MultispeQ at 35 DPI (n=10). Means with no letter in common are significantly different (Tukey's multiple comparison test; $p<0.05$). Errors bars indicate ± SEM.
