## Supplemental Figure 2 for "Integrated Systems-Level Proteomics and Metabolomics Reveals the Diel Molecular Landscape of Diverse Kale Cultivars"

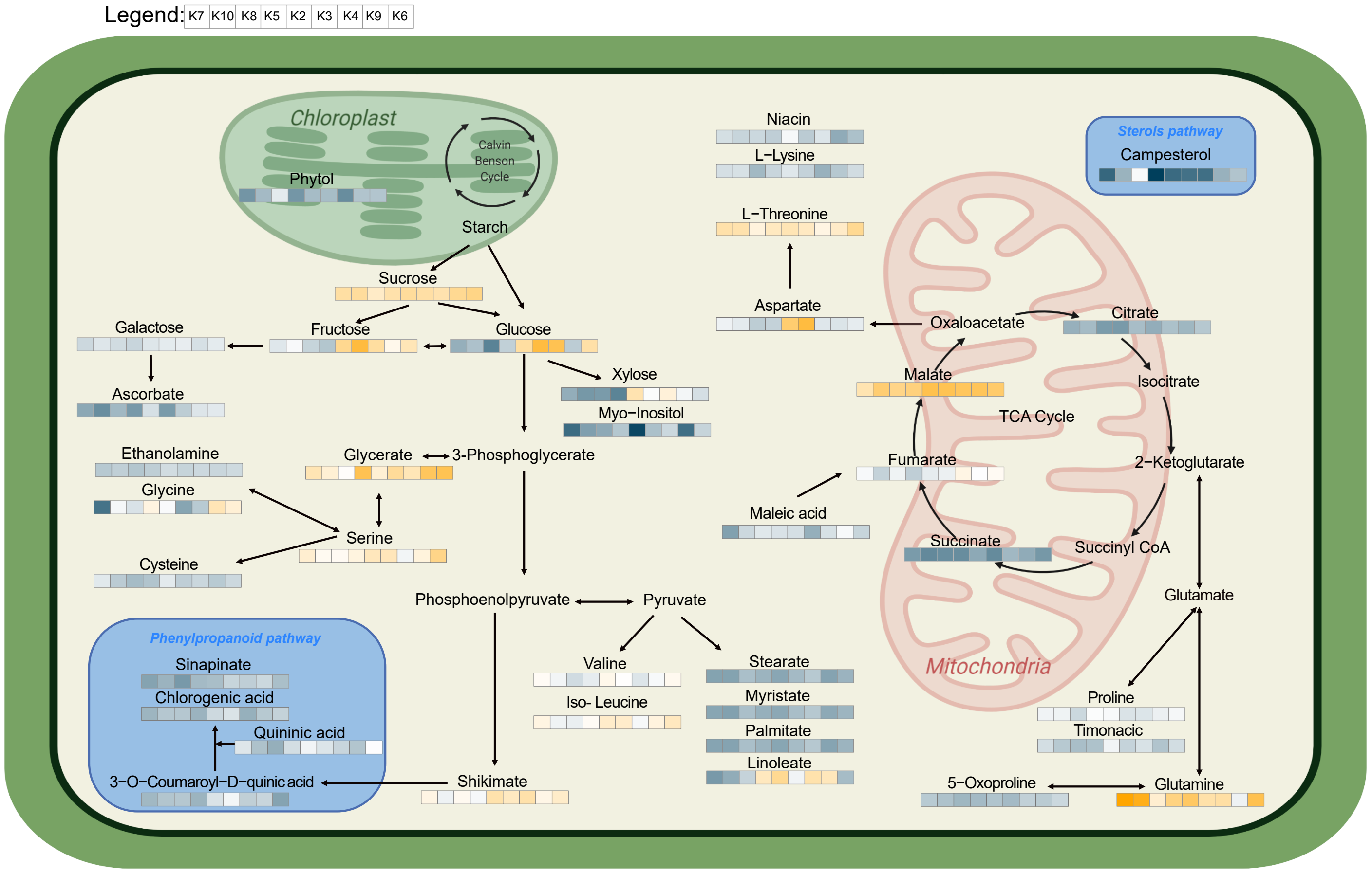


**Supplemental Figure 2: Diel changes in the kale metabolic landscape.** Simplified schematic of mapped diel metabolic changes observed in kale cultivars by GC-MS.
